# Structural basis of bulk lipid transfer by bridge-like lipid transfer protein LPD-3

**DOI:** 10.1101/2024.06.21.600134

**Authors:** Yunsik Kang, Katherine S. Lehmann, Juan Vanegas, Hannah Long, Amanda Jefferson, Marc Freeman, Sarah Clark

**Affiliations:** Vollum Institute, Oregon Health & Science University, Portland, Oregon 97239, USA; Department of Biochemistry and Biophysics, Oregon State University, Corvallis, Oregon 97331, USA

## Abstract

Bridge-like lipid transport proteins (BLTPs) are an evolutionarily conserved family of proteins that localize to membrane contact sites and are thought to mediate the bulk transfer of lipids from a donor membrane, typically the endoplasmic reticulum (ER), to an acceptor membrane, such as a that of the cell or an organelle ^1^. Despite the fundamental importance of BLTPs for cellular function, the architecture, composition, and lipid transfer mechanisms remain poorly characterized. Here, we present the subunit composition and the cryo-electron microscopy structure of the native LPD-3 BLTP complex isolated from transgenic *C. elegans*. LPD-3 folds into an elongated, rod-shaped tunnel whose interior is filled with ordered lipid molecules that are coordinated by a track of ionizable residues that line one side of the tunnel. LPD-3 forms a complex with two previously uncharacterized proteins, here named “Intake” and “Spigot”, both of which interact with the N-terminal end of LPD-3 where lipids enter the tunnel. Intake has three transmembrane helices, one of which borders the entrance to the tunnel; Spigot has one transmembrane helix and extends 80 Å along the cytosolic surface of LPD-3. Experiments in multiple model systems indicate that Spigot plays a conserved role in ER-PM contact site formation. Our LPD-3 complex structural data, together with molecular dynamics simulations of the transmembrane region in a lipid bilayer, reveal protein-lipid interactions that suggest a model for how the native LPD-3-complex mediates bulk lipid transport and provide a foundation for mechanistic studies of BLTPs.

## Introduction

Cells and cellular organelles are encircled by membranes that each contain a characteristic lipid composition ^2^. These membranes are not static structures, but are constantly undergoing dynamic modification as a result of cellular processes like autophagy, phagocytosis, exocytosis, organelle biosynthesis or even changing environmental conditions (e.g. temperature). The majority of cellular lipids are synthesized in the endoplasmic reticulum (ER) and, since lipids cannot efficiently diffuse through the aqueous cytosolic environment, they are delivered to other cellular compartments via vesicular or nonvesicular transport ^3, 4^. Nonvesicular lipid transport is carried out by lipid transport proteins (LTPs), cytoplasmic proteins that contain a hydrophobic core which shelters lipids from the aqueous environment during transport ^5^. LTPs are primarily found at membrane contact sites, such as those between the ER and the plasma membrane, and most known LTPs operate by shuttling single lipid molecules one at a time from the donor to acceptor membrane. In contrast, the recently discovered bridge-like lipid transport proteins (BLTPs) are believed to function as molecular bridges that span the distance between the ER and membranes targeted for lipid delivery, enabling the solubilization and transport of large volumes of lipids ^1, 6^. Structural evidence ^7–9^ and Alphafold predictions ^1^ indicate that BLTPs fold into a rod-like tunnel with a hydrophobic lining that is formed by a series of repeating β-groove domains (RBG domains), a defining feature of the BLTP superfamily.

Mutations in human BLTP1 are linked to Alkuraya-Kučinska syndrome, a devastating neurological disorder ^10^, but the cellular and molecular basis of this syndrome is not understood. BLTP1 family members are found in all eukaryotes and have been demonstrated to localize to ER-plasma membrane (PM) contact sites in both yeast and worms ^11, 12^. Multiple lines of evidence support the notion that BLTP1 and its orthologs function as molecular bridges for lipid transport at ER-PM contact sites: in yeast, the BLTP1ortholog (Csf1) is important for growth at low temperatures and homeoviscous membrane adaptation ^11, 13, 14^; in *Drosophila*, the BLTP1 ortholog (Tweek) is required for development, synaptic vesicle recycling ^15^, astrocyte-mediated phagocytosis of neuronal debris ^16^; and in *C. elegans*, the BLTP1 ortholog (LPD-3) is critical for animal development, efficient phagocytosis, and adaptation to cold stress^12, 16^.

While it is believed that the primary function of BLTP1 family members is bulk transfer of lipids from the ER to the plasma membrane, the mechanisms of lipid uptake and transfer, regulation of lipid transport, and potential additional roles for BLTPs *in vivo* remain undefined. All BLTP1 orthologs possess a very high molecular weight – LPD-3 and Tweek are the 32^nd^ and 23^rd^ largest proteins in the worm and fly genomes, respectively – and their enormous size has hindered structural and biochemical analysis due to difficulties associated with recombinant production of full-length protein ^17^. In this study, we overcome these challenges by isolating the native LPD-3 complex from transgenic *C. elegans*. Single particle cryo-EM reconstructions and mass spectrometry analyses reveal the subunit composition and structural details of the native LPD-3 complex, including its interaction with 27 lipid molecules that fill the hydrophobic LPD-3 tunnel. Combining this structural information with molecular dynamics simulations, we present a model for the structural basis of LPD-3-mediated lipid transfer between membranes.

## Results

### The native LPD-3 complex includes two previously uncharacterized proteins

To enable isolation of the native LPD-3 complex, we generated two transgenic worm lines in which DNA encoding an mVenus fluorophore and a 3XFLAG tag were inserted either at the start or the end of the endogenous *lpd-3* gene. We refer to these as the *mvenus::lpd3* and *lpd3::mvenus* worm lines, which respectively produce N- and C-terminally tagged LPD-3 protein. To determine if insertion of the tag at either position disrupts LPD-3 protein function in the transgenic worms, we compared the development rates and cold tolerance of the transgenic worm lines to that of wildtype N2 worms. We found that the C-terminally tagged *lpd3::mvenus* worms and WT worms developed at approximately the same rate, with more than 90% of both reaching the L4 developmental stage by 40 h (Extended Data Figure 1a). The *lpd3::mvenus* worms and WT worms also exhibited a similar resilience to cold exposure, with ∼80% of both worm populations surviving after 12 hours at 4°C, consistent with previous results ^12^ (Extended Data Figure 1b). By contrast, only 14% of N-terminally tagged *mvenus::lpd3* worms reached the L4 stage by 40 h and only 35% survived cold exposure, indicating impaired LPD-3 function. The C-terminally tagged *lpd3::mvenus* worm line was therefore selected for downstream structural and biochemical experiments. Further characterization of the *lpd3::mvenus* worm line by spectral confocal imaging revealed a pattern of mVenus fluorescence that was consistent with previously reported images of tagged-LPD-3 ^12^ (Extended Data Figure 2a).

Next, using our recently developed protocol for isolating native membrane protein complexes from *C. elegans* ^18^, the C-terminally tagged LPD-3 complex was isolated from *lpd3::mvenus* worms (Extended Data Figure 2). During purification, fluorescence-based size exclusion chromatography (FSEC) gave a monodisperse peak with an estimated molecular weight of 750 kDa (Extended Data Figure 2b). Since this value exceeds the expected 483 kDa size of tagged LPD-3, this implies that the complex harbors additional subunits.

To determine the subunit composition of the LPD-3 complex, we performed mass spectrometry (MS) analyses (Extended Data Figure 2d). MS results contained peptides that spanned the entire 4022-residue LPD-3 protein, indicating that the isolated protein was intact. In addition to the LPD-3 peptides, two proteins stood out in terms of their high peptide spectral counts and appeared to have co-purified with LPD-3. Based on their locations in the cryo-EM structure (see below), we have named them Spigot and Intake. These are both previously uncharacterized proteins, although the human ortholog of Spigot, C1orf43, was identified along with BLTP1 in a genome-wide CRISPR screen as a regulator of phagocytosis of *Legionella pneumophila* in macrophages ^19^. Neither Spigot nor Intake was observed in an identically prepared sample from wildtype N2 worms, demonstrating that these proteins specifically associate with LPD-3.

Sequence alignment searches revealed that Spigot is widely conserved across the animal kingdom and exhibits a level of sequence conservation roughly matching that of LPD-3 with its orthologs. For both proteins, the *C. elegans, D. melanogaster* and human orthologs exhibit 15-20% sequence identity. Interestingly, Spigot and its orthologs are predicted to be ER-resident proteins, as assessed by the program ERPred ^20^. In contrast, sequence searches for Intake yielded no recognizable orthologs outside of the nematode family.

Further inspection of the MS analyses revealed twelve additional proteins that exhibit much lower peptide spectral counts compared to Spigot and Intake yet show a strong contrast with the control sample from wildtype worms (Extended Data Figure 3a). To investigate the possibility that these proteins functionally interact with LPD-3 complex *in vivo* (e.g. transiently or by binding to a subset), we assayed the ability of astrocytes to phagocytose neuronal debris during developmental neuronal remodeling, a recently discovered role for *Drosophila* Tweek (Tweek is the LPD-3 ortholog) ^16^. We found that astrocyte-specific RNAi knockdown of 9 of the 12 additional proteins that co-purified with the LPD-3 complex resulted in significant defects in astrocyte phagocytic function (Extended Data Figure 3). These data support the notion that the majority of these co-purified proteins are also required for BLTP1 complex function *in vivo*.

### Structure determination and overall architecture of the LPD-3 complex

To elucidate the architecture of the native, C-terminally-tagged LPD-3 complex, we performed single molecule cryo-EM on the purified complex. We initially obtained a map of the complex at a nominal resolution of 2.7 Å that corresponds to the N-terminal half of LPD-3 (Extended Data Figures 4 and 5 and Extended Data Table 1). In an effort to resolve the entire 4022 residue protein, we selected particles with strong density features for the C-terminal portion of the protein and obtained a ‘full-length’ map at a nominal resolution of 6.2 Å (Extended Data Figure 6). We suspect that the inability to obtain a high-resolution map of the C-terminal half of LPD-3, residues 1680-4022, is due to conformational heterogeneity in that half of the protein. Notably, the starting point of the heterogeneity matches the location of a 115-residue segment – residues 1680-1794 – that is predicted to be intrinsically disordered, and the C-terminal half of LPD-3 is predicted to contain 8 loops, from 37 to 71 residues long, that are also predicted to be disordered. As the LPD-3 complex is presumably anchored to the membrane at the N- and C-terminus, this conformational heterogeneity may confer necessary structural flexibility.

The ‘full-length’ map measures ∼345 Å at its longest axis, which matches the expected size of LPD-3 based on Alphafold2 predictions (Extended Data Figure 6). Density corresponding to a detergent micelle is observed at one end of the ‘full-length’ map, indicating the presence of a transmembrane domain, and a tunnel is observed along the length of the entire protein (Extended Data Figure 6b). These features are consistent with predicted BLTP1 models that contain a single N-terminal transmembrane helix ^1^, as well as with low resolution cryo-EM maps of BLTP family members Vps13 ^7^, SHIP164 ^8^, and Atg2 ^21^.

The ‘N-terminal’ map allowed us to build a model of LPD-3 residues 21-1679, Spigot residues 7- 187, and Intake residues 208-303, along with 30 well-resolved lipid molecules (Figure 1). LPD-3 begins with a single transmembrane helix, followed by an elongated, tube-shaped structure with an internal cavity. The protein fold is consistent with experimental structures of recombinantly-expressed fragments ^7–9, 22^ and predicted models ^1, 23^ of other BLTP family members. The LPD-3 tunnel is positioned at a ∼40° angle relative to the membrane bilayer and is filled with an abundance of well-resolved phospholipids. Spigot extends ∼70 Å across the exterior surface of LPD-3, beginning with its transmembrane helix and terminating with an irregular four-helix bundle (helices H2 to H5) that is nested on the surface of LPD-3 (Figure 1). As Spigot is a predicted ER-resident protein, its location reinforces existing evidence ^12^ that LPD-3 is N-terminally anchored to the ER. Given that assumption, this structure likely represents the site where lipids enter the tunnel for transport to the plasma membrane. The visible portion of Intake is entirely within the micelle, forming a three-helix transmembrane bundle that, along with Spigot, helps anchor LPD-3 in the membrane. Intake transmembrane helix 2 (TM2), along with the LPD-3 TM1, create a surface within the membrane that leads to the entrance to the lipid transport tunnel (Figure 1b). The Intake and Spigot subunits interact minimally with each other, as their transmembrane helices are located on opposite sides of the LPD-3 TM1.

**Figure 1.**
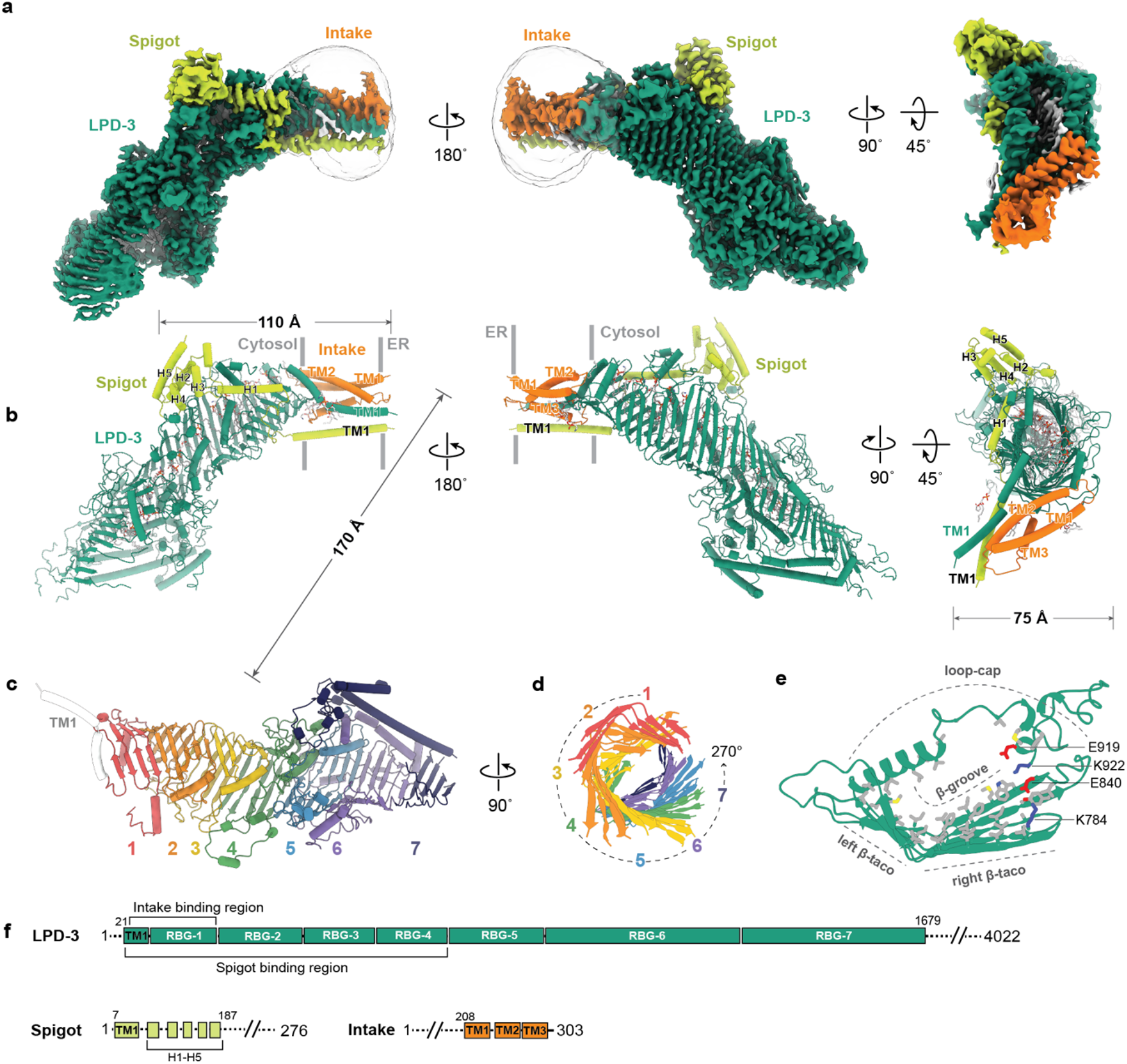
Architecture of the LPD-3 complex. **a,** Cryo-EM map of the native LPD-3 complex is shown parallel to the membrane and perpendicular to the membrane. LPD-3 is colored teal, Spigot is colored yellow, Intake is colored orange, and lipid molecules are colored light grey. **b,** Model of the native LPD-3 complex, shown in the same views and colors as (a). **c,** LPD-3 is colored according to RBG domain and TM1 is colored white. Spigot and Intake are hidden for clarity. The numbers corresponding to each RBG domain are shown at the bottom. **d,** Top-down view of LPD-3, highlighting the spiraling RBG architecture. Only the β-taco of each RBG domain is shown and RBG domains are colored as in (c). **e,** Cartoon representation of RBG-5, showing the composition of tunnel-lining residues. Hydrophobic residues are colored grey, positively charged residues are colored blue, and negatively charged residues are colored red. Each element of the RBG architecture is labeled. **f,** Schematic representation of LPD-3, Spigot, and Intake. TM, transmembrane domain. H, helix. Dashed lines indicate regions of the protein that were not visible in the cryo-EM map.

### Structural features of LPD-3 and lipid organization within the tunnel

The LPD-3 model includes 7 of its 16 RBG domains ^1^, which are stacked in tandem to form the rod-shaped hydrophobic tunnel that extends into the cytosol from the N-terminal transmembrane helix. Each modeled RBG domain consists of 4-5 anti-parallel ß-strands that are arranged in a ß**-**taco fold, followed by a ‘loop-cap’ structure that is comprised of extended coil regions with one or more α-helices (Figure 1c). The loop-cap covers the ß**-**taco groove and links adjacent RBG domains. The length of the loop-cap ranges from 36 residues in RBG-3 to 64 residues in RBG-5. Multiple RBG domains also harbor long segments between ß-strands, the longest of which is a 226-residue segment in RBG-6 that meanders along the outer surface of the ß**-**taco, associating mainly via hydrophobic interactions. The width of tunnel gradually increases from ∼25 Å in RBG-1 to ∼40 Å in RBG-7 (roughly the tunnel midpoint), due to a combination of flattening of the ß**-**taco and lengthening of the RBG β-strands. The sequential RBG domains form a regular, spiraling pattern involving a rotation of ∼38° per domain (Figure 1d). RBG-1 and RBG-7 are therefore offset by nearly 270°. As RBG domains are a defining feature of the BLTP superfamily, our structure provides a valuable experimental view of this unique architecture and demonstrates that the tunnel gets larger as one moves away from the entrance.

The amino acid composition of the tunnel interior creates an environment that is ideal for solubilization and transport of membrane phospholipids. Hydrophobic residues line the ß-groove and the interior-facing surfaces of the loop-caps (Figure 1e). Additionally, a striking string of ionizable residues form a track that extends along the right edge of the ß-taco and the bottom of the loop-cap (Figure 1e). This track runs the full length of the tunnel from the membrane to RBG-7 (Extended Data Figure 7). Based on the lipid orientations in the structure, these largely alternating acidic and basic residues are well-positioned to coordinate the head groups of the phospholipids and facilitate their transport through the tunnel.

We observed 30 density features modeled as full and partial lipid molecules within the transmembrane region, the tunnel entrance, and the tunnel (Figure 2). Three phospholipid molecules decorate the transmembrane helices, one of which, PL1, is buried in a cavity formed by transmembrane helices from all three proteins and appears to play a structural role (Figure 2a). The entrance to the tunnel is bordered by LPD-3 TM1 and Intake TM2, which form a V-shaped track with the opening of the V at the tunnel entrance, potentially creating a path to shuttle lipids from the membrane environment into the hydrophobic interior of the tunnel (Figure 2a). At the tunnel entrance, one partial lipid is modeled that forms hydrophobic interactions with Intake residues F245 and F247 (Figure 2b). Within the tunnel, a single continuous row of phospholipids are lined up edge-to-edge that span the full length of the tunnel (Figure 2c, Extended Data Figure 5g). Some parts of the tunnel are larger and accommodate additional rows of lipids that are stacked up to five deep (Figure 2d). The lipid orientations are such that all of the headgroups are near the ionizable residues along the right edges of the β-taco folds and the acyl tails extend towards the opposing edge (Figure 2f). A similar orientation was seen for the single ordered phospholipid observed in the crystal structure of the Atg2 N-terminal RBG domain ^9^.

**Figure 2.**
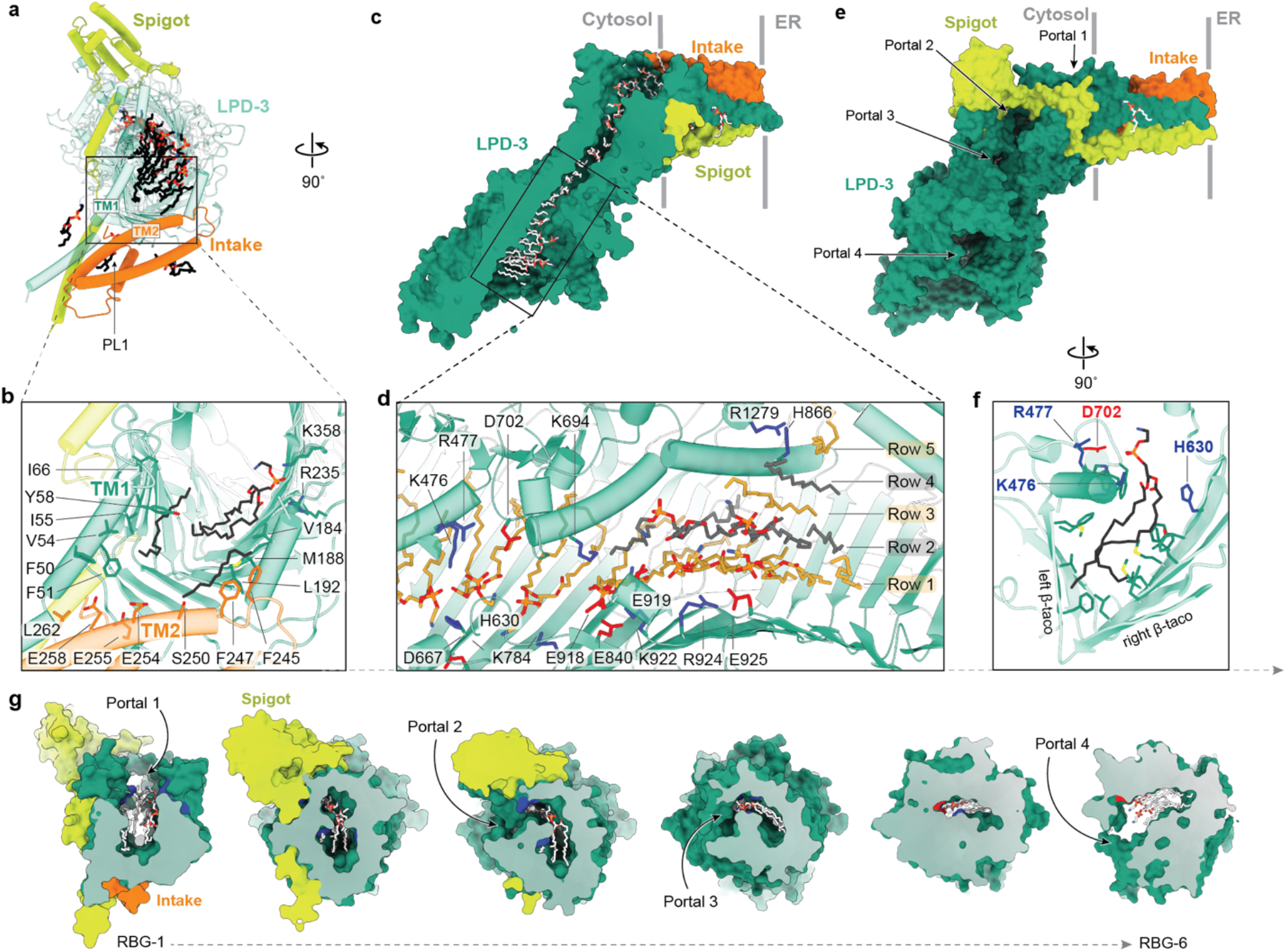
Organization of lipids in the LPD-3 tunnel. **a,** Overview of the lipids observed in the LPD-3 complex, shown perpendicular to the membrane. Proteins are shown in cartoon representation. LPD-3 is colored teal, Spigot is colored yellow, and Intake is colored orange. Lipids are shown as black sticks. **b,** Zoomed in view of the tunnel entrance, colored as in (a). **c,** Clipped view of the cryo-EM map to show the lipid transport tunnel. LPD-3, Spigot, and Intake subunits are shown in surface representation. Proteins are colored as in (a) and lipids are shown as white sticks. **d,** Organization of lipids within the tunnel. Ionizable residues that interact with lipid head groups are labeled and shown as sticks. Lipids are shown as grey or gold sticks. **e,** The LPD-3 complex viewed parallel to the membrane. Arrows indicate the location of the four hydration portals. The same colors are used as in (a). **f,** Zoomed view of a lipid within the tunnel. Residues that interact with the lipid headgroup are shown as sticks. **g,** Sliced views of the LPD-3 tunnel to show the locations of the hydration portals. Slices are sequential from left to right and progress from RBG-1 to RBG-6.

Unexpectedly, the LPD-3 tunnel is punctuated by four openings that expose the interior to the aqueous cytosol (Figure 2e). These ‘hydration portals’ are located in the loop-caps along the right side of the β-taco, in line with the charged residues that form the ionizable track. The portals, which range in width from 6-8 Å at their narrowest point, are ideally positioned to enable water molecules to access the head groups of the phospholipids (Figure 2g). We speculate that hydration of phospholipid head groups facilitates lipid diffusion through the tunnel by mimicking the environmental conditions of a lipid bilayer; the acyl tails are enclosed in a hydrophobic environment and the head groups are exposed to the solvent. These conditions likely lower the energetic costs associated with lipid transport compared to a sealed tunnel.

The number of stacked rows in various parts of the tunnel depends on the tunnel diameter. In RBG-1 and RBG-2, the tunnel diameter is sufficient to accommodate two rows of lipids. The tunnel narrows in RBG-3 and RBG-4 to fit a single row of lipids and then gradually expands through RBG-5 and RBG-6 to hold 2 to 5 rows. The increase in tunnel width and lipid rows is accompanied by an increase in the number of ionizable residues (Extended Data Figure 7). While only seven ionizable residues line the interior of RBG-1 and RBG-2, eleven such residues are observed in RBG-5 and RBG-6, presumably to coordinate a larger number of lipid headgroups. The interior of RBG-7 has room for lipids but no density to guide their modeling. There are an average of 4.5 lipid molecules per RBG domain within the tunnel, which means that the full-length LPD-3 molecule with 16 RBG domains can theoretically accommodate 72 lipids. Consistent with this estimation, molecular modeling of BLTPs suggest that Further, we note that the packing density of the lipids within the tunnel is similar to that of phospholipids in a modeled membrane bilayer; the average distance between phosphate head groups is 8.4 Å within the tunnel and 9 Å within a membrane bilayer that is modeled for molecular dynamics simulations (see section ‘Intake borders the entrance of the lipid transfer tunnel’ below). Bulk lipid transfer therefore resembles small patches of a membrane leaflet moving through a tunnel.

### Spigot forms extensive interactions with LPD-3 and is critical for ER-PM contact site formation

Spigot orthologs are found throughout the animal kingdom, from nematodes to humans. Spigot makes extensive contacts with LPD-3, starting in the membrane and wrapping along the tunnel exterior, burying ∼3,600 Å^2^ of LPD-3 surface (Figure 3). The Spigot transmembrane helix (TM1) engages with LPD-3 TM1 through hydrophobic interactions, as well as through a hydrogen bond between Spigot Arg31, which is highly conserved (Figure 3e), and the LPD-3 Leu41 backbone carbonyl oxygen (Figure 3b). A linker domain that includes H1 connects Spigot TM1 to a four-helix bundle. A mixture of electrostatic and hydrophobic interactions anchor the linker domain to the outer surface of LPD-3 RBG-1 and RBG-2 (Figure 3c). The four-helix bundle is perched on the cytosolic surfaces of both RBG-3 and RBG-4, where the tunnel opening ends and the narrowest part of the tunnel begins. The bundle is oriented nearly perpendicular to the LPD-3 tunnel. The bundle docks into a cleft formed by a protruding loop from the RBG-4 β-taco fold, making multiple electrostatic and hydrophobic interactions (Figure 3d). The Spigot residues that interact with LPD-3 tend to be well conserved (Figure 3e), implying that Spigot is a conserved subunit of functional LPD-3 complexes.

**Figure 3.**
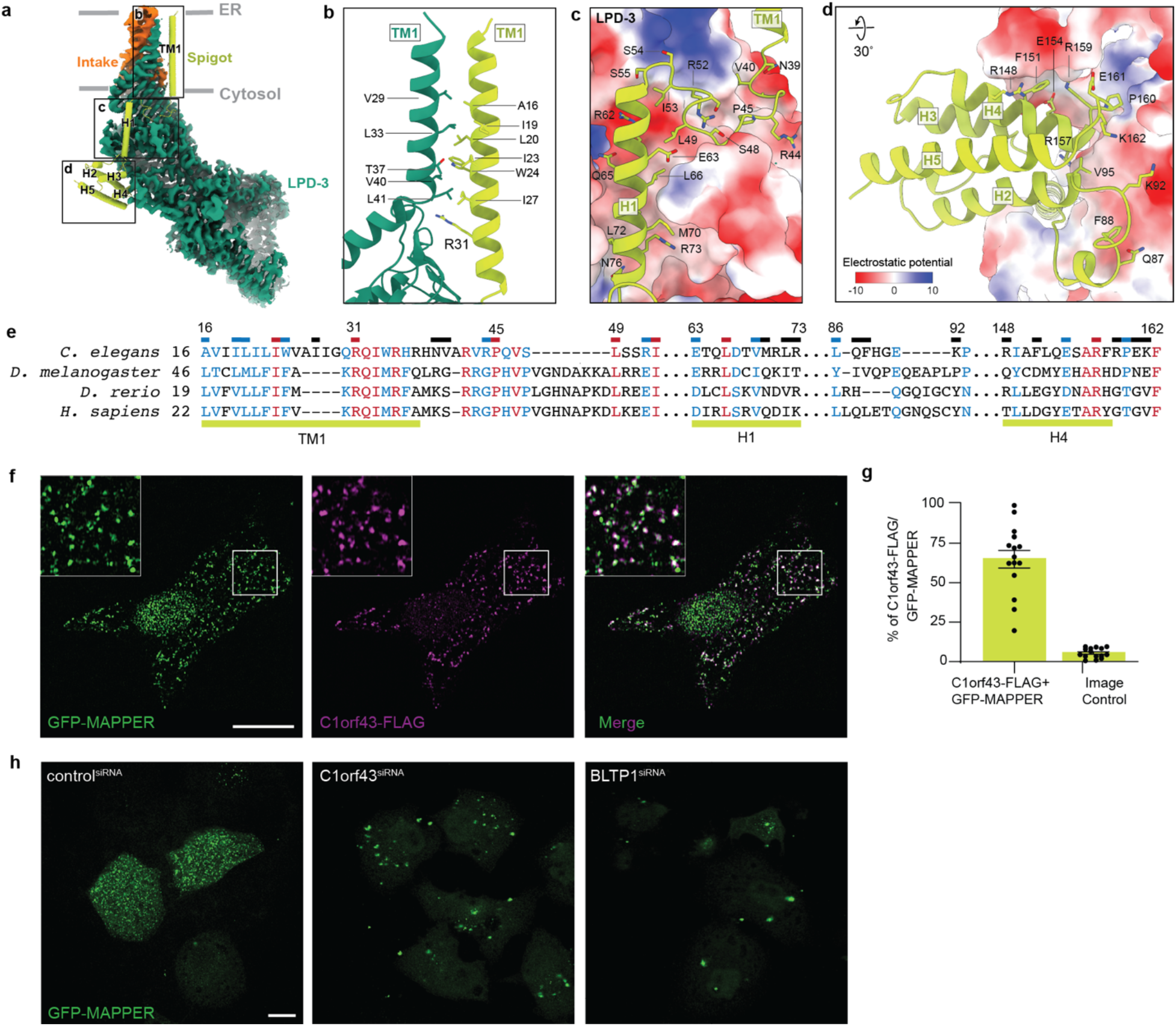
Spigot interactions with LPD-3 and localization in human cell culture. **a,** Binding interface between Spigot and LPD-3. Cryo-EM map is shown for LPD-3 and Intake. Spigot is shown as a cartoon. **b,** The interface between LPD-3 TM1 and Spigot TM1. Interacting residues are shown as sticks. **c, d,** The interface between LPD-3 and Spigot H1 (**c**) and LPD-3 and Spigot H2-H5 (**d**). The electrostatic surface of LPD-3 is shown, where blue represents positive regions and red represents negative regions. Spigot is shown in yellow. **e,** Multiple sequence alignment of Spigot orthologues. Elements of secondary structure are shown below the sequences and residues that interact with LPD-3 are indicated above with horizontal bars. Residues in black are not conserved, those in blue are conservatively substituted, and those in red are conserved, as assessed by the program PROMALS3D ^45^. **f, g,** Confocal microscopy imaging of HeLa cells co-expressing GFP::Mapper to mark ER-PM contact sites and C1orf43::FLAG (**f**) reveal significant overlap upon quantification (**g**). **h,** Knockdown of endogenous C1orf43 or BLTP1 with siRNA leads to ER-PM contact site collapse as visualized by GFP::Mapper.

To investigate whether Spigot is important for ER-PM contact site formation, we first examined the localization of recombinant C1orf43, the human ortholog of Spigot, in HeLa cells. When HeLa cells were co-transfected with plasmids encoding C1orf43::FLAG and GFP::MAPPER, a GFP reporter that selectively labels ER-PM contact sites ^24^, we observed robust localization of C1orf43 at ER-PM junctions (Figure 3f, g). Next, we used siRNA to knockdown endogenously expressed *C1orf43* in HeLa cells, which resulted in a collapse of ER-PM junctions that mirrors the *BLTP1* knockdown phenotype (Figure 3h), indicating that C1orf43 is important for ER-PM contact site stability. The *Drosophila* BLTP1 ortholog, Tweek, is similarly required for proper formation of ER-PM contact sites ^16^. These results, together with our data showing that Spigot (CG6665 in flies) is important for astrocyte-mediated phagocytosis in flies (Extended Data Figure 3), are consistent with a conserved role for Spigot in BLTP1- mediated lipid transfer functions at ER-PM contact sites.

### Intake borders the entrance of the lipid transfer tunnel

We observed density for the C-terminal 95 residues of Intake in the cryo-EM map, which fold into three transmembrane helices (TM1, TM2 and TM3) (Figure 1f). Our model is consistent with the Alphafold predicted secondary structure, which shows Intake residues 1-180 as largely disordered and residues 181- 303 as forming four alpha helices, the last three of which were modeled in our structure. There is additional cryo-EM density on the ER leaflet side of the membrane that is that is consistent with a few turns of a low resolution helix (Figure 1a), which possibly corresponds to the fourth alpha helix. The Intake three-helix bundle docks with LPD-3 at the tunnel entrance, burying ∼1,800 Å^2^ surface area. The interaction between Intake and LPD-3 is mediated primarily by the short, cytosolic loop that links Intake TM1 and TM2 (Figure 4b). Hydrophobic interactions between the LPD-3 transmembrane helix and Intake TM1 strengthen the complex. The C-terminal 8-residues of Intake are well-ordered and its C-terminal carboxylate interacts with a cluster of arginine residues, two of which are from LPD-3 and one of which is from Spigot (Figure 4c), solidifying the association of all three subunits. A buried phospholipid molecule (PL1) that is enclosed in a cavity formed by transmembrane helices from all three subunits appears to further stabilize the complex (Figure 4c).

**Figure 4.**
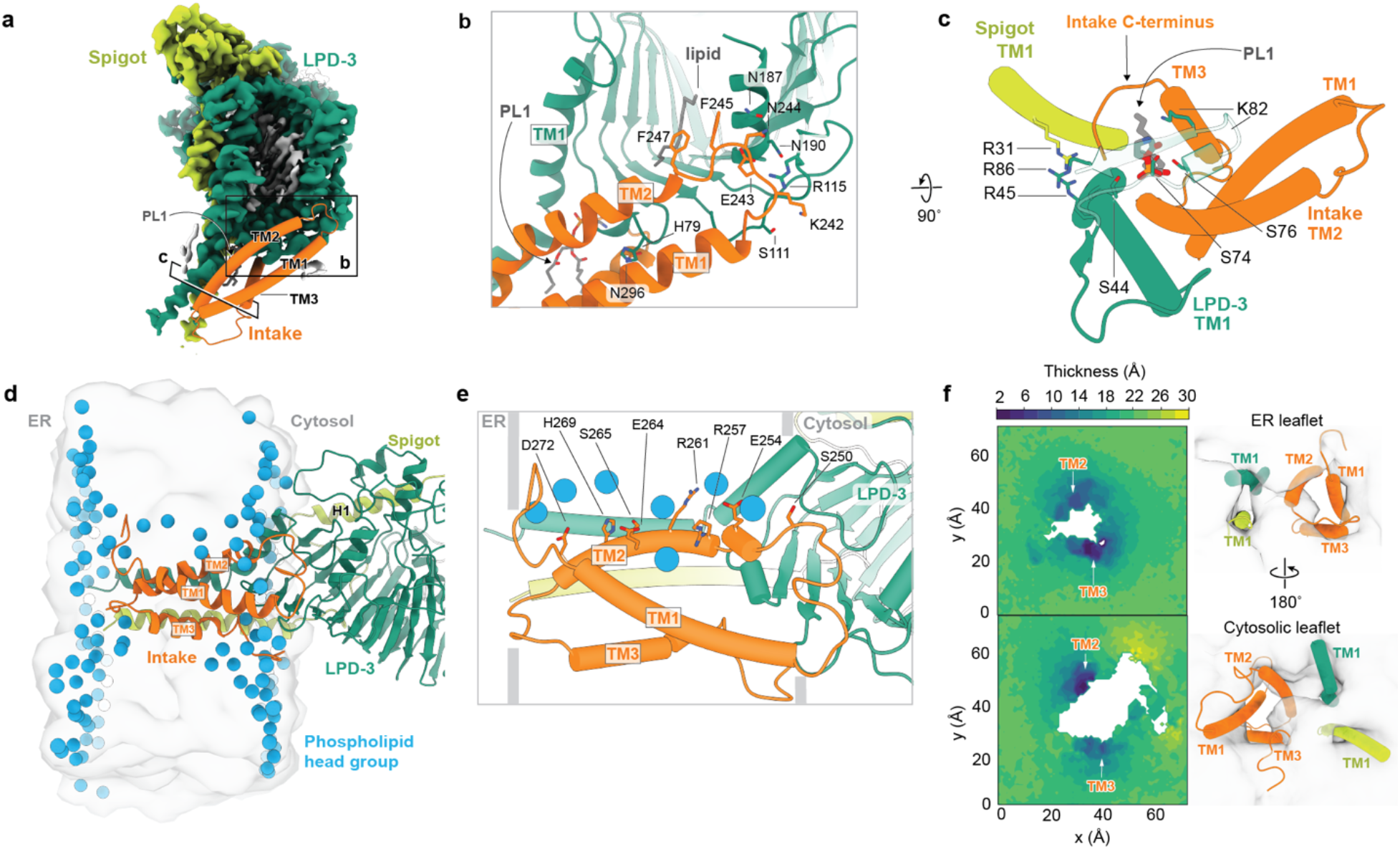
Intake interactions with LPD-3 and membrane integration of the complex. **a**, The LPD-3 complex viewed perpendicular to the membrane. Intake is shown in cartoon representation and is colored orange. Cryo-EM density for LPD-3 is colored teal, Spigot is colored yellow, and lipids are colored grey. **b,** Interaction of Intake with LPD-3. Lipid molecules are shown as grey sticks and interacting residues are shown as sticks. **c,** The LPD-3 complex viewed perpendicular to the membrane. Interacting residues are shown as sticks and phospholipid PL1 is colored grey. **d**, Molecular dynamics simulation of the membrane-embedded LPD-3 complex shows lipids distributed along the interior of the bilayer. Phosphate atoms are shown as blue spheres and the membrane is shown as a transparent grey surface. Proteins are colored as in (a). **e**, Interactions between Intake TM2 and lipid molecules. Interacting residues are shown as sticks and phosphate molecules are shown as blue spheres. **f**, Average leaflet thickness in the vicinity of the protein complex as measured by the position of the phosphate headgroups relative to the bilayer midplane. Data averaged over the last 250 ns of the trajectory with the system is re-centered around the protein for each simulation frame analyzed. The cross section of the protein is white and the membrane is colored according to thickness. Cartoon representations of the same view of the LPD-3 complex are shown to the right of each thickness plot.

Interestingly, all three of the Intake TM helices harbor an unusually large number of polar and ionizable residues that are not buried in the protein interior, but rather are exposed to the non-polar part of the membrane, a feature that we hypothesize has important functional consequences for lipid entry into LPD-3. Specifically, TM2 features eight membrane-exposed ionizable and polar residues, while TM1 and TM3 harbor six and three such residues, respectively. Although pairs of salt bridge-forming ionizable residues have been observed in transmembrane helices in other proteins ^25^, we were unable to find an example of a protein that harbors so many polar residues within their transmembrane domains, highlighting the unusual composition of these helices.

To understand how Intake alters the lipid bilayer and interacts with individual lipids, we performed all-atom molecular dynamics simulations on the membrane-proximal part of the LPD-3 complex embedded in a membrane composed of phospholipids and cholesterol. To stabilize the Intake TM3 helix during the simulations, it was necessary to place a phospholipid in the location of PL1 (Figure 4c), consistent with a structural role for this lipid. The most notable interactions occur with TM2, where a line of phospholipids traverse the bilayer with the head groups of the phospholipids interacting with Intake TM2 residues Arg257, Arg261, Glu264, Thr268, and Asp272, and the tails oriented nearly perpendicular to the bilayer (Figure 4d-e). Inspection of the equilibrated membrane reveals a strong perturbation of the membrane structure around Intake TM2 and TM3 and analysis of membrane leaflet thickness shows that both the cytoplasmic and ER leaflets are markedly thinner adjacent to TM2 and TM3 (Figure 4f).

Based on this remarkable line up of phospholipids traversing the membrane, we speculate that Intake TM2 functions like a track along which lipids are shuttled from the ER leaflet of the membrane to the tunnel. In support of this hypothesis, the polar Intake TM2 residues that interact with phospholipid molecules in our simulations are located on the same side of the tunnel as the track of LPD-3 ionizable residues that coordinate headgroups. Taken together, the charged residues in Intake TM2 and the polar and solvent-exposed parts of the LPD-3 tunnel form a continuous track for the phospholipid headgroups that leads from the lumen of the ER through the tunnel and presumably ends at the plasma membrane.

### Mechanistic insights into LPD-3-mediated lipid transport

Accumulating evidence from *in vivo* and *in vitro* studies indicate that BLTPs function as molecular bridges for lipid transport at ER-membrane contact sites. Our work reveals the subunit identity and architecture of the native LPD-3 complex, as well as key protein-lipid interactions that illuminate the organization and flow of lipids from the ER membrane, into the tunnel, and along the tunnel interior.

The ‘core’ complex is composed of the rod-shaped LPD-3 protein and the auxiliary proteins Spigot and Intake. Spigot interacts extensively with N-terminus of LPD-3 and is essential for proper ER-PM contact site formation in HeLa cells (Figure 3h) and astrocyte-mediated phagocytosis in flies (Extended Data Figure 3), suggesting that it is a conserved subunit of BLTP1 complexes. While the role of Spigot in ER-PM contact formation and LPD-3-mediated lipid transfer is unclear, it may stabilize LPD-3 or aid in localizing the N-terminus to the ER. Intake resides in the membrane near the entrance of the LPD-3 tunnel and appears to function as a ‘lipid escalator’ that ferries lipids from the ER and into the tunnel. MD simulations indicate that the unusually high number of polar amino acids in Intake’s transmembrane helices interact with phospholipid head groups to both destabilize the membrane and move lipids to the tunnel. Intriguingly, there is no ortholog for Intake outside of the nematode family, raising the question of which protein performs an analogous function in other organisms.

In addition to the core subunits Spigot and Intake, it is likely that other, transiently associated auxiliary proteins interact with the LPD-3 complex to regulate lipid transfer and promote localization of the N- and C-termini to the ER and plasma membrane, respectively. Through a combination of mass spectrometry and RNAi screening, we identified nine proteins that are important for astrocyte-mediated phagocytosis (Extended Data Figure 3), but further studies are required to determine whether they are involved in LPD-3-mediated lipid transfer. It is possible that some of these proteins aid in localizing the LPD-3 C-terminus to the plasma membrane, a process for which the mechanism is currently unknown due to the lack of a C-terminal transmembrane helix. The BLTP family members Vps13 and Atg2, which also lack a transmembrane helix at their C-terminus, associate with the donor membrane via interaction with binding partners ^26, 27^.

Based on our structure and MD simulations, we propose a model of LPD-3-mediated bulk lipid transport. Phospholipids are shuttled from the ER membrane and into the tunnel via interaction of the head groups with polar amino acids in Intake TM2. The lipids are then ‘handed off’ to ionizable residues that line the interior of LPD-3 on the right side of the β-taco. A series of hydration portals are interspersed with the ionizable residues that allow water molecules to enter the tunnel and hydrate the lipid head groups. Together, these hydration portals and alternately charged ionizable residues create a continuous track that facilitates lipid flow and prevents stagnation within the tunnel. Although we do not see the C-terminal half of the LPD-3 tunnel due to presumed conformational heterogeneity, we assume that this track continues to the plasma membrane. Future experiments of full-length LPD-3 and its orthologs will be required to fully elaborate the mechanism of lipid transport and uncover additional elements of the process, such as lipid selectivity and integration of the tunnel endpoint in the donor membrane. Finally, as mutations in human BLTP1 have been implicated in AKS ^10, 28^, our studies establish a framework for structure-based investigations of the disease.

## Materials and Methods

### Transgenic worm design

The strain *PHX5024 lpd-3 (syb5024)* was generated by SunyBiotech using CRISPR/Cas9 genome editing and is referred to as the *lpd3::mVenus* line (Extended Data Figure 1). The mVenus-3xFLAG sequence was inserted prior to the stop codon of the endogenous *lpd-3* gene (Wormbase: Y47G6A.23). The genotype was confirmed using PCR and primers ER16-seq-s (GAAAAGTACAATCCCTGCCA) and ER16-seq-a (ACTTTCTGCGTTTGCGTGTG), which bind 469 bp upstream and 318 bp downstream from the insertion site, respectively to amplify the region of interest. The strain *PHX5131 lpd-3 (syb5131)* was generated by SunyBiotech using CRISPR/Cas9 genome editing and is referred to as the *mVenus*::*lpd3* line (Extended Data Figure 1). The mVenus-3xFLAG sequence was inserted prior to the start codon of the endogenous *lpd-3* gene. The genotype was confirmed using PCR and primers ER15-seq-s (AAATGCACAAAACCTGTGTG) and ER15-seq-a (AAGCGATCAAAGAGGAGAAT), which bind 432 bp upstream and 388 bp downstream from the insertion site, respectively to amplify the region of interest. To enable elution of the engineered LPD-3 complex from affinity chromatography resin, a PreScission protease (3C) cleavage site was placed between the mVenus fluorophore and 3xFLAG tag for both worm lines.

### Spectral confocal imaging

Adult and larval worms were immobilized in M9 buffer (22 mM KH_2_PO_4_, 42 mM Na_2_HPO_4_, 86 mM NaCl, and 1 mM MgCl_2_) containing 30 mM sodium azide and placed on slides that were prepared with ∼4 mm agar pads. Spectral images were acquired on a Zeiss 34-channel LSM 880 Fast Airyscan inverted microscope with a 40x 1.2 NA water-immersion objective lens. Linear unmixing was employed to distinguish between the mVenus signal and autofluorescence. The autofluorescence signal was subtracted from each image. The 3D z-stack information is presented in 2D after performing a maximum intensity projection.

### C. elegans development and cold resilience assays

Developmental and cold resilience assays were performed in accordance with protocol developed by Wang et al ^12^. Worms were cultured at 20°C on HB101 seeded nematode growth media (NGM) for at least four generations and were never starved. For both developmental and cold resilience assays, worms were bleach synchronized and plated on NGM of equal thickness seeded with equal amounts of HB101.

To analyze the developmental rate of wildtype and tagged LPD-3 worm lines, bleach synchronized worms were cultured at 20°C on NGM. After 40-, 45-, or 50-hours post-plating, the percentage of worms reaching the L4 stage (determined by the presence of a crescent-shaped vulva) were counted.

To analyze the cold resilience of wildtype and tagged LPD-3 worm lines, bleach synchronized worms were cultured for 50 hours at 20°C on NGM. At 50 hours, worms were counted and the presence of any dead recorded. Immediately following, worms were placed at 4°C for 20 hours, then removed and allowed to recover at 20°C for 24 hours before the percentage of survivors was determined. Worms were considered dead if they appeared necrotic, displayed no pharynx pumping, and did not move in response to plate tapping when observed using light microscopy. Statistical analyses were performed using GraphPad Prism 9.

### Large scale *C. elegans* culture

Large-scale worm growth and optimization of LPD-3 protein isolation conditions were performed as described in Clark et al ^18^. Briefly, nematode growth medium (NGM) agar plates were prepared and spread with *E. coli* strain HB101, allowing the bacterial lawn to grow overnight at 37 °C. Worms were transferred to the NGM plates and grown for 3-4 days at 20° C until HB101 cells were depleted. Worms on the plates were transferred to a liquid medium in 2 L baffled flasks, supplemented with HB101 (∼15 g per 500 mL medium) and streptomycin (50 μg/mL), and worms were grown at 20° C with vigorous shaking (150 rpm) for 70-72 hours. To harvest worms, the liquid culture flasks were placed on ice for 1 hour to allow the worms to settle. The media was removed and the worm slurry was collected in a tube, washed twice with 50 mL of ice cold M9 buffer by successive centrifugation (800 x g for 1 minute) and resuspension. Worms were ‘cleaned’ by sucrose density centrifugation at 1500 x g for 5 minutes after bringing the volume of worm slurry up to 25 mL with M9 buffer and adding 25 mL of ice cold 60% (w/v) sucrose. The worm layer on top was recovered and placed in a new tube and then washed twice with 50 mL of ice cold M9 buffer. The volume of the worm pellet was measured and the same volume of M9 buffer was added to the tube and worm balls were made by dripping the slurry into liquid nitrogen. The worm balls were stored at −80 °C until further use.

### Isolation of the native LPD-3 complex

Approximately 40 g of frozen worm balls were disrupted using a ball mill (MM400, Retch) where the grinding jar and ball were pre-cooled in liquid nitrogen. Disrupted worm powder was solubilized at 4 °C for 2.5 hours in a buffer containing 50 mM Tris-Cl (pH 8), 150 mM NaCl, 5 mM EDTA, 2% (w/v) glyco-diosgenin (GDN), and protease inhibitors (0.8 µM aprotinin, 2 µg/ml leupeptin and 2 µM pepstatin). After centrifugation at 40,000 rpm (186,000 x g) for 50 minutes, the supernatant was filtered through a 0.45 µm filter and incubated with anti-FLAG M2 affinity for 2.5 hours on a rotator at 4°C. The resin was washed 5 times with a buffer containing 20 mM Tris-Cl (pH 8), 150 mM NaCl and 0.02% (w/v) GDN, using a volume of buffer that was 200-fold the volume of the resin. The LPD-3 complex was eluted by incubating with 10 µg of 3C protease at 4°C for 1 hour on the rotator. The sample was concentrated to 100 µL using a 100 kDa MWCO concentrator and the concentrate was loaded onto a size-exclusion chromatography (SEC) column (Superose 6 Increase 10/30 GL, GE Healthcare), equilibrated in a buffer composed of 20 mM Tris-Cl (pH 8), 150 mM NaCl, 0.02% (w/v) GDN. The peak fractions from the putative LPD-3 complex were pooled and concentrated for cryo-EM grid preparation or mass spectrometry experiments. Approximately 250 ng of LPD-3 was isolated from 40 g of worm balls, which are equal to 3×10^7^ worms. The amount of protein was determined via mVenus fluorescence based on comparison to an mVenus standard curve on the FSEC. The isolated native LPD-3 sample was analyzed by SDS-PAGE (sodium dodecyl sulfate–polyacrylamide gel electrophoresis) and the protein bands were visualized by silver staining.

The same isolation method was utilized to make the wild-type worm sample from the *C. elegans* N2 strain for use as a control in the mass spectrometry experiments. Approximately 40 g of N2 worm balls were purified using anti-FLAG M2 resin, followed by purification on an SEC column, and fractions in the same elution volume as LPD-3 were collected, pooled and concentrated. The concentrate was subjected to mass spectrometry analysis in order to evaluate non-specific binding of *C. elegans* proteins to anti-FLAG M2 affinity resin.

### Mass spectrometry

The SEC-purified LPD-3 complex sample was dried and then dissolved in a solution consisting of 5% sodium dodecyl sulfate, 8 M urea, and 100 mM glycine (pH 7.55). The sample was then reduced with [tris(2-carboxyethyl)phosphine (TCEP)] at 37 °C for 15 min and alkylated with methyl methanethiosulfonate for 15 min at room temperature, followed by addition of acidified 90% methanol and 100 mM triethylammonium bicarbonate buffer (TEAB; pH 7.55). The sample was then digested in an S-trap micro column briefly with 2 µg of a Tryp/LysC protease mixture, followed by a wash and 2-h digestion at 47 °C with trypsin. The peptides were eluted with a solution consisting of 50 mM TEAB, 50% acetonitrile and 0.2% formic acid, and were then pooled and dried. Each sample was dissolved in 20 µL of 5% formic acid and injected into the Thermo Fisher QExactive HF mass spectrometer. Protein digests were separated using a Dionex RSLC UHPLC system, then delivered to a QExactive HF (Thermo Fisher) via electrospray ionization using a Nano Flex Ion Spray Source (Thermo Fisher) that is fitted with a 20-μm stainless steel nano-bore emitter spray tip and a 2.6 kV source voltage. Xcalibur version 4.0 was used to control the system. Samples were applied at 10 µL/min to a Symmetry C18 trap cartridge (Waters) for 10 min, then switched onto a 75 µm × 250 mm NanoAcquity BEH 130 C18 column with 1.7 µm particles (Waters) using mobile phases water (A) and acetonitrile (B) containing 0.1% formic acid and a 7.5 to 30% acetonitrile gradient over 60 min with a 300 nL/min flow rate. Survey mass spectra were acquired over m/z 375 to 1,400 at 120,000 resolution (m/z 200), and data-dependent acquisition selected the top 10 most abundant precursor ions for tandem mass spectrometry by higher energy collisional dissociation using an isolation width of 1.2 m/z, normalized collision energy of 30, and a resolution of 30,000. Dynamic exclusion was set to auto, charge state for MS/MS +2 to +7, maximum ion time 100 ms, minimum AGC target of 3 × 106 in MS1 mode, and 5 × 103 in MS2 mode. Data analysis was performed using Comet (v. 2016.01, rev. 3)^29^ against a January 2022 version of canonical FASTA protein database containing C. elegans uniprot sequences and concatenated sequence-reversed entries to estimate error thresholds and 179 common contaminant sequences and their reversed forms. Comet searches for all samples were performed with trypsin enzyme specificity. Monoisotopic parent ion mass tolerance was set to 1.25 Da and monoisotopic fragment ion mass tolerance was set to 1.0005 Da. A static modification of +45.9877 Da was added to all cysteine residues and a variable modification of +15.9949 Da on methionine residues. A linear discriminant transformation was used to improve the identification sensitivity from the Comet analysis ^30, 31^. Separate histograms were created for matches to forward sequences and for matches to reversed sequences for all peptides of seven amino acids or longer. The score histograms of reversed matches were used to estimate peptide false discovery rates (FDRs) and set score thresholds for each peptide class. The overall protein FDR was 1.2%.

### *Drosophila* RNAi screen

Flies (*Drosophila melanogaster*) were reared on standard cornmeal agar under a 12-hour light/12-hour dark cycle at 25°C. All experimental procedures were carried out at 25°C. Auxiliary binding partners of LPD-3 were screened, focusing on proteins unrelated to transcription, translation, protein folding, or the cytoskeleton. *Drosophila* homologs of these selected proteins were subjected to RNAi-mediated knockdown in astrocytes using the *GMR25H07-Gal4* driver line (Bloomington, 49145). For each RNAi condition, five animals were dissected at head eversion (HE), a developmental stage occurring approximately 12 hours into metamorphosis, characterized by the emergence of the immature adult head.

The following lines were used: *UAS-FLP5.DD*, *UAS- tweek* RNAi (VDRC 110686), *CG6665* RNAi (Bloomington 60402), *Pepck2* RNAi (VDRC 107092), *Apoltp* RNAi (Bloomington 51937), *CG33303* RNAi (VDRC 107778), *FASN1* RNAi (VDRC 108339), *SERCA* RNAi (VDRC 107446), *SdhA* RNAi (VDRC 110440), *Rack1* RNAi (VDRC 104470), *PCB* RNAi (VDRC 105936), *Rab1* RNAi (Bloomington 27299), *Idh* RNAi (VDRC 100554).

For sample collection, prepupae were placed on a 1% agarose plate, and the dissection of the central nervous system (CNS) was conducted every 10 minutes at HE. Dissected samples were initially placed in PBS with 0.1% Triton X-100 (0.1% PBST) on ice and then transferred to a solution containing 4% formaldehyde (PFA) in PBST for fixation, which lasted for 20 minutes at room temperature (RT) with gentle agitation. Following fixation, samples underwent two washes in 0.1% PBST and were subsequently incubated with primary antibodies in 0.1% PBST on a shaker at 4°C overnight. Washes were conducted in 0.1% PBST at RT for five minutes each, followed by overnight incubation at 4°C on a shaker in 0.1% PBST. Afterwards, samples were exposed to secondary antibody in 0.1% PBST for 3 hours at room temperature, followed by additional washing and storage in 0.1% PBST overnight at 4°C. Finally, samples were mounted using Vectashield antifade reagent. The primary antibody used for larvae staining was α-Crz (Crz, 1:1000), and the secondary antibody used was Cy3 (Jackson ImmunoResearch). Fixed fly CNS samples were imaged using a Zeiss LSM 880 confocal microscope. Image tiles were stitched together. Area of vCrz neurites (without cell bodies) where quantified and statistics were performed using GraphPad Prism 9.

### Cell Culture

HeLa cells were maintained at 37 °C in Dulbecco’s Modified Eagle Medium (DMEM) (Gibco) with 10% FBS and 4 mM glutamine (complete medium, CM). Cells were cultured on chambered cover glass (18mm diameter, #1 ½ thickness cover glass, Electron Microscopy Sciences). Cells that were transfected with a plasmid alone were transfected with GFP-Mapper (Addgene 117721) or C1ORF43-FLAG (SinoBiologic) constructs using lipofectamine 3000 (Invitrogen), according to the manufacturer’s instructions. Cells were transfected at the 100 ng-200 ng range. Cells were fixed and imaged 48 hours following transfection. Cells that were transfected with plasmid and siRNA were transfected with DharmaFECT Duo Transfection Reagent according to the manufacturer’s instructions. SMARTPools of siRNA were used for *KIAA1109* (*BLTP1*) and *C1orf43*. Cells were fixed and imaged 48 hours following transfection of siRNA.

All experiments were performed in 12-well cell culture plates (TrueLine) and once cells were transfected they were maintained in Opti-MEM Reduced Serum Medium (Gibco). Cells were fixed in 0.1% Triton X-100 (0.1% PBST) with 1% formaldehyde for 30 minutes, washed, put in blocking buffer (1% BSA in PBS) for 30 minutes and then washed again. Primary antibody stains were done overnight at 4 °C on a shaker. Primary was then removed, and the samples were washed three times with PBS. Secondary was applied for 1 hour at room temperature while shaking. Samples were washed with PBS and mounted on slides with Prolong Gold antifade (Invitrogen). The primary antibodies used for cell culture stain were as follows: : α-GFP (abcam 1:1000),and α-FLAG (Sigma-Aldrich 1:250).

Samples were imaged using a Zeiss LSM 880 confocal microscope with Airyscan Fast mode. For colocalization analysis, a 100 μm² region of cells was used to calculate the percentage of C1orf43-FLAG expression in GFP::MAPPER. For control images, each GFP::MAPPER image was horizontally flipped, and colocalization was calculated. Statistical analyses were performed using GraphPad Prism 9.

### Cryo-EM sample preparation

A volume of 3.5 μL of the concentrated LPD-3 complex was applied to a Quantifoil grid (R2/1 300 gold mesh, covered by 2 nm continuous carbon film), which was glow-discharged at 15 mA for 30 seconds in the presence of amylamine. The grids were blotted and flash frozen using a Vitrobot mark IV for 2.5 seconds with 0 blot force after 10 seconds wait time under 100% humidity at 10°C. The grids were plunge-frozen into liquid ethane, cooled by liquid nitrogen.

### Data acquisition

The native LPD-3 complex dataset was collected on a 300 keV FEI Titan Krios microscope equipped with a K3 detector. The micrographs were acquired in super-resolution mode (0.394 Å/pixel) with a magnification of 29 Kx corresponding to a physical pixel size of 0.788 Å/pixel. Images were collected by a 3×3 multi-hole per stage shift and a 7 multi-shot per hole method using Serial EM, with a defocus range of −1.5 to -2.5 μm. Each movie stack consisted of 50 frames per movie, with a total dose of 50 e-/ Å^2^. A total of 8,195 movies were collected.

### Image processing

Beam-induced motion was corrected by patch motion correction with an output Fourier cropping factor of 1/2 (0.788 Å/pixel). Contrast transfer function (CTF) parameters were estimated by patch CTF estimation in CryoSparc v3.3.1. A total of 8,061 movies were selected by manual curation and the particles were picked using blob-picker with an elliptical blob in which the minimum and maximum particle diameters were set to 100 and 400 Å, respectively. Initially, 3.9 million particles were picked and, following particle inspection, 2.6 million particles were extracted with a box size of 756 pixels, 4x binned (3.152 Å/pixel). After three rounds of 2D classification, ‘junk’ particles were removed, resulting in 938,732 million particles in total. The classes with the highest resolution features were selected to generate an *ab initio* model. All 938,732 particles were then subjected to one round of heterogenous refinement using the *ab initio* model as a reference to further remove ‘junk’ particles. The best class contained 861,293 particles that were re-extracted from unbinned micrographs and subjected to one more round of heterogeneous refinement. Two classes, containing 713,216 particles total, were selected for non-uniform refinement and the resulting map was used to generate templates for template picking. Approximately 2.1 million particles were extracted from unbinned micrographs with a box size of 756 pixels and subjected to two rounds of 2D classification and one round of heterogeneous refinement. The best class, which contained 285,377 particles, was selected for non-uniform refinement, yielding a 2.7 Å reconstruction of the N-terminal half of LPD-3.

We noted that many of our 2D classes contained features that are consistent with a full-length protein. We selected the classes with the strongest full-length features and re-extracted them with a box size of 1024, 2x binned (1.576 Å/pixel). All 146,403 particles were used to generate an *ab initio* model, which was then subjected to non-uniform refinement. As the refined volume was centered near the N-terminus, we used the volume alignment tool in CryoSparc to re-center the volume on the middle of the protein. Non-uniform refinement of the shifted volume resulted in a nominal 6.2 Å reconstruction of the full-length LPD-3 complex.

### Atomic model building and refinement

The “N-terminal” density map was sharpened with Phenix AutoSharpen, and both sharpened and unsharpened maps were used for structure determination. The initial model of LPD-3, residues 1-1950, was predicted by Alphafold2 ^32^ and fit into the map with rigid body fitting in UCSF ChimeraX ^33^, followed by manual adjustment and *de novo* model building in Coot v0.9.8.92 ^34^. *De novo* regions were built by tracing the backbone, assigning the sequence, and fitting the side chains using Coot. The model was refined with the real space refinement program in PHENIX v1.21 ^35^. Although there is density corresponding to one more RBG domain (RBG-8) in the map, the domain was not modeled due to low resolution. The following regions of LPD-3 were not modeled due to lack of density: 1-20, 164-180, 1115-1146, 1209-1231, 1299-1301, 1327-1339, 1355-1380, 1400-1420, and 1680-4022.

The structure of the Spigot auxiliary protein was built using a combination of *de novo* model building and rigid body fitting of the Alphafold2 predicted structure. Each α-helix was individually docked into the map and manually adjusted in Coot, using the side chains of bulky tryptophan, tyrosine, and phenylalanine residues, including W24, Y94, F151 and F163, as guides to correctly assign register. Spigot residues 1-6 and 188-276 were not modeled due to lack of density.

The structure of the Intake auxiliary protein was built manually into the density map using Coot. The well-resolved C-terminus of the protein, which is located in a cavity of the transmembrane region, was used as a guidepost to facilitate the definition of the correct register of the Intake sequence. A pair of phenylalanine residues, F245 and F247, that reside near the entrance of the LPD-3 tunnel further helped to assign the sequence register. Intake residues 1-209 were not modeled due to lack of density, however, an additional density feature was observed in the cryo-EM map that extends away from Intake TM1, towards the ER lumen side of the micelle, and is roughly shaped like an alpha helix (Figure 1). The shape and location of this density feature suggests that it corresponds to the C-terminal end of a soluble helix, which is consistent with the Alphfold2 predicted structure of Intake residues 172-195. The density feature is too ill-defined for model building.

The phospholipid molecules were identified based on a ‘V’ shaped density feature and all phospholipids were modeled as 1,2-dioleoyl-sn-glycero-3-phosphoethanolamine (PEE). The number of carbon molecules in the acyl chain were modified based on the density; the length of acyl chains ranged from C3 to C16. Partial lipid densities were built in regions where an acyl chain of 10 or more carbons could be fit in the density feature.

### Molecular dynamics simulations

The GROMACS simulation package ^36^ v2023 was used to perform molecular dynamics (MD) simulations. Proteins and lipids were represented using the CHARMM36m force-field (FF) ^37–39^ with the default CHARMM-modified TIP3P water model, which includes additional VdW parameter on the H atoms. The combined protein assembly was modified in the following ways: the LPD-3 model was truncated to include residues 21 to 405, the Spigot model was truncated to include residues 7 to 76, and a disordered loop that was not visible in the cryo-EM map was modeled for LPD-3 residues 164-180. The protein assembly was then processed through the PPM webserver (https://opm.phar.umich.edu/ppm_server) to obtain a model with the correct orientation and position relative to the lipid membrane ^40^. The oriented protein complex was uploaded to the CHARMM-GUI webserver (https://www.charmm-gui.org/) to build the combined membrane-protein system ^41, 42^. All protein residues were simulated in their default protonation states at neutral pH except for Glu292, which is buried within the protein and was simulated in the neutral protonated state. The membrane composition was chosen to roughly mimic the endoplasmic reticulum and includes the following lipids: 87 POPE (1- palmitoyl-2-oleoyl-glycero-3-phosphoethanolamine), 51 POPC (1-palmitoyl-2-oleoyl-glycero-3- phosphocholine), 12 PSM (palmitoylsphingomyelin), and 10 cholesterol. The upper and lower leaflets have a slight difference in the number of lipids to accommodate the shape of the transmembrane region of the protein. Sufficient water was added to have a layer of at least 1 nm next to the protein and potassium and chloride ions were added to neutralize the system and have concentration corresponding to 150 mM KCl. Electrostatic interactions were computed using the particle-mesh Ewald method with a real space cutoff 1.2 nm and a Fourier grid spacing of 0.12 nm, while Van der Waals interactions were computed using a force-switched Lennard-Jones potential between 1.0 and 1.2 nm. Newton’s equations of motion were integrated with a leap-frog algorithm using a 2 fs time step. The temperature of the system was held constant at 37 °C using a velocity-rescaling algorithm ^43^ with a time constant of 1 ps, and the pressure was held constant at 1 atm using a semi-isotropic stochastic cell-rescaling algorithm ^44^ using a time constant of 5 ps. Three independent replicas were run with different initial membrane configurations. Each replica was first energy minimized for 1,000 steps using a steepest descent algorithm, followed by a 250 ns membrane equilibration period were the protein was under harmonic position restraints using spring constants of 1,000 kJ/mol,nm^2^ for the protein backbone and 500 kJ/mol,nm^2^ for the sidechains. After equilibration, each replica was simulated without any restraints for 2.5 microseconds. Positions of all atoms were saved to the trajectory in 100 ps intervals.

## Acknowledgements

We thank P. Andrew Karplus for help with structure analysis and manuscript preparation. We thank Eric Gouaux and members of the Gouaux lab for helpful discussions. We thank Ashok Reddy for mass spectrometric analysis, Farzad Jalali-Yazdi for help with cryo-EM data processing, Sepehr Dehghani for scripts for calculating membrane thickness, and Marzia Miletto for help with cryo-EM screening and data collection. Cryo-EM grid screening and data collection were performed at the Pacific Northwest Cryo-EM Center (PNCC), which is supported by NIH grant U24GM129547. The OHSU Proteomics Shared Resource is partially supported by NIH core grants P30EY010572 and P30CA069533. This work was supported by HHMI funding to E.G., Oregon State University startup funds to S.C., NIH grant K99 NS126642 to Y.K., NIH grants R37 NS053538-18 and R01 NS124146-01 to M.F. and NSF grant CHE- 1944892/2326678 to J.V.

## Author Contributions

S.C. performed the cryo-EM experiments. Y.K. and A.J. performed the RNAi screen and astrocyte-mediated phagocytosis assays. Y.K. and K.S.L. performed the cell culture experiments. H.L performed functional analyses of the tagged worm lines. J.V. performed and analyzed MD simulations. Y.K., M.F, and S.C. designed the project and wrote the manuscript. All authors contributed to manuscript preparation.

## Data Availability

The volumes for the cryo-EM data have been deposited in the Electron Microscopy Data Bank under accession codes EMD-45276 (full-length) and EMD-45399 (N-terminal). The coordinates of the LPD-3 complex have been deposited in the Protein Data Bank under accession code 9CAP.

## Materials & Correspondence

Correspondence and material requests should be addressed to Sarah Clark.

---

**Extended Data Figure 1.**
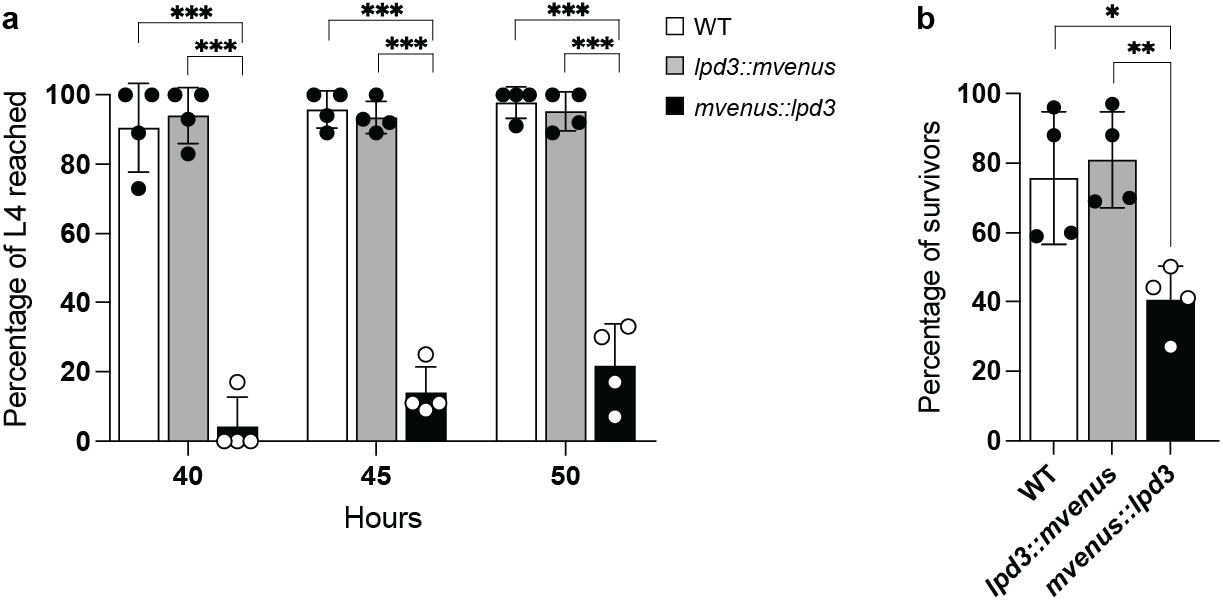
Development and cold tolerance of wild type and transgenic worms. **a,** Percentages of animals that reached developmental L4 stage at indicated hours post egg preparation in wild type (WT) worms, *lpd3::mvenus* worms, or *mvenus::lpd3* worms. Values are means +/- S.D. ***P < 0.001 (two-sided unpaired t-test, N = 4 biological replicates) **b,** Percentages of animals that survived cold exposure (4 °C for 20 hours) post egg preparation in wild type (WT) worms, *lpd3::mvenus* worms, or *mvenus::lpd3* worms. *P<0.05, **P<0.01 Values are means +/- S.D. (two-sided unpaired t-test, N = 4 biological replicates).

**Extended Data Figure 2.**
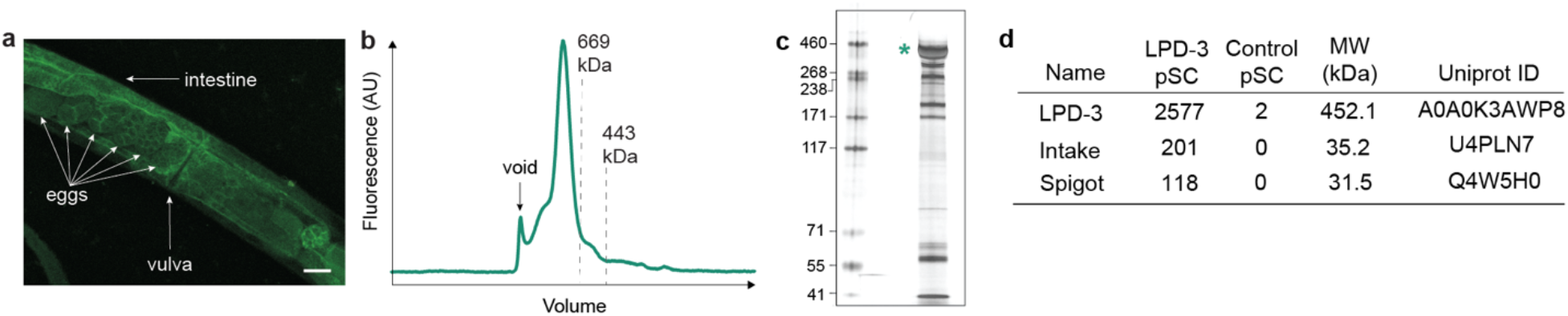
Isolation of LPD-3 from *C. elegans*. **a,** Spectral confocal image of mVenus fluorescence in an *lpd3::mvenus* worm showing fluorescence throughout the worm body. **b,** Representative FSEC profile of the LPD-3 complex, detected via tryptophan fluorescence. **c,** A silver stained SDS-PAGE gel of the purified LPD-3 complex. Teal asterisk indicates LPD-3. Other bands may correspond to transiently associated proteins or contaminants. **d,** Mass spectrometry analysis of the purified LPD-3 complex, showing the three proteins with the highest peptide spectral counts (pSC).

**Extended Data Figure 3.**
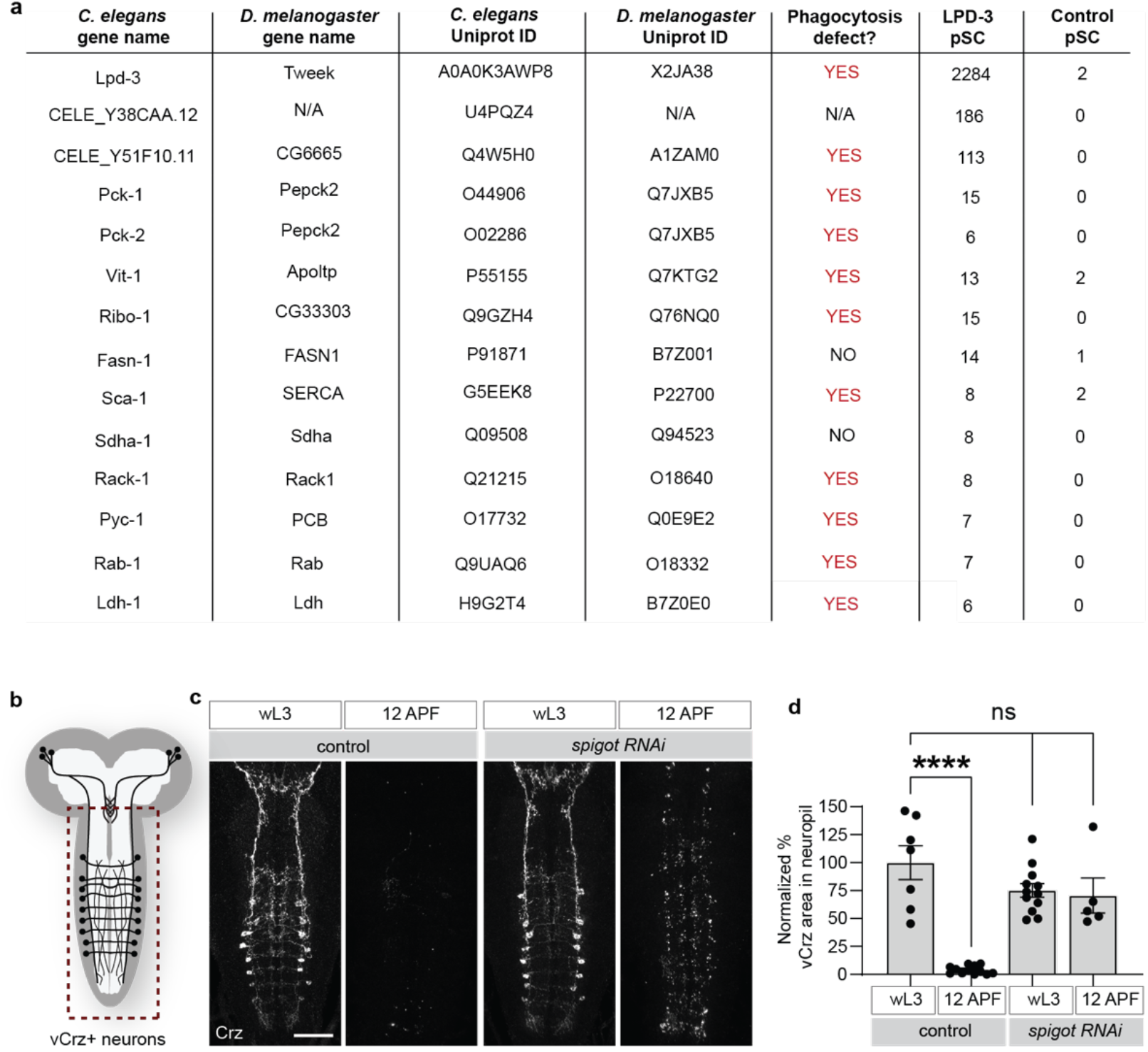
RNAi screen in *Drosophila*. **a,** Mass spectrometry analysis of the LPD-3 complex, showing the proteins that were subjected to RNAi-mediated knockdown in *Drosophila* astrocytes to in order to identify regulators of phagocytosis. pSC = peptide spectral counts. **b,** Overview of vCrz neuron morphology: Eight pairs of vCrz neurons are present in the ventral nerve cord, red dotted box represents imaged region. **c**, vCrz neuronsat the wandering third instar larval stage (wL3) in controls which is broken down and fully cleared by astrocytes at head eversion (HE, ∼12 hours after puparium formation) in controls. Astrocyte knockdown of *spigot* (*CG6665*) with *GMR25H07-Gal4* (astrocyte-specific driver) does not affect neuronal morphology at wL3 but disrupts the ability of astrocytes to clear neuronal debris by HE. vCrz neurons were labeled with anti-Crz, and all images are maximum Z projections of the entire ventral nerve cord (VNC). Scale bar, 50 μm. **d**, Quantification of data from (**c**) for controls (*UAS-FLP5.DD*) (wL3, N=7; HE, N=12) and *UAS-spigot RNAi* (wL3, N=12; HE, N=5). Graphs show mean ± SEM, and each dot represents independent animals. Statistical comparisons were performed using two-way ANOVA with Sidak’s multiple comparisons test. ****p<0.0001, ns=not significant.

**Extended Data Figure 4.**
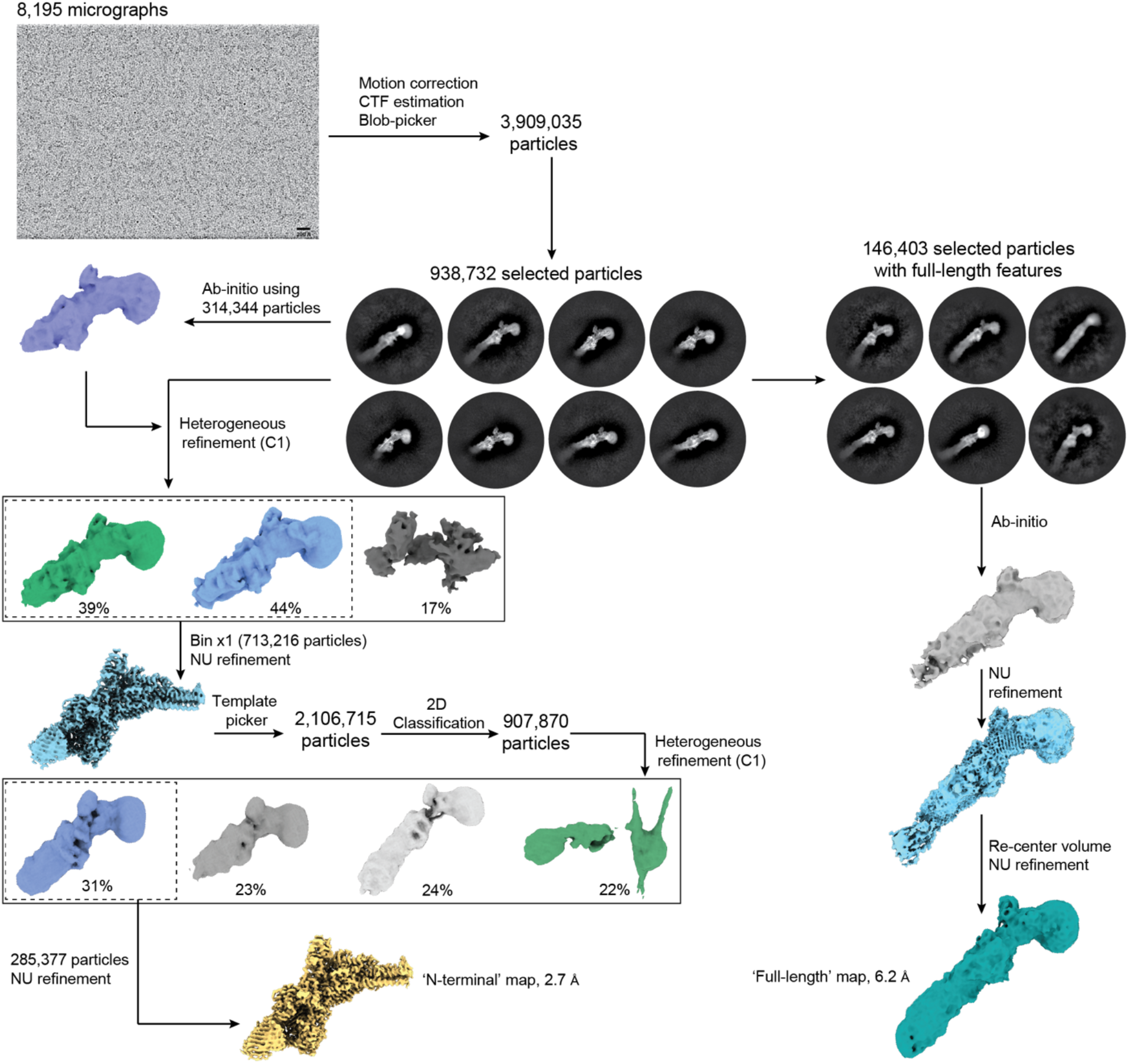
Cryo-EM data analysis of the LPD-3 complex. Flow chart for cryo-EM data analysis of the LPD-3 “N-terminal” map and “full-length” map. Scale bar of cryo-EM micrograph = 200 Å.

**Extended Data Figure 5.**
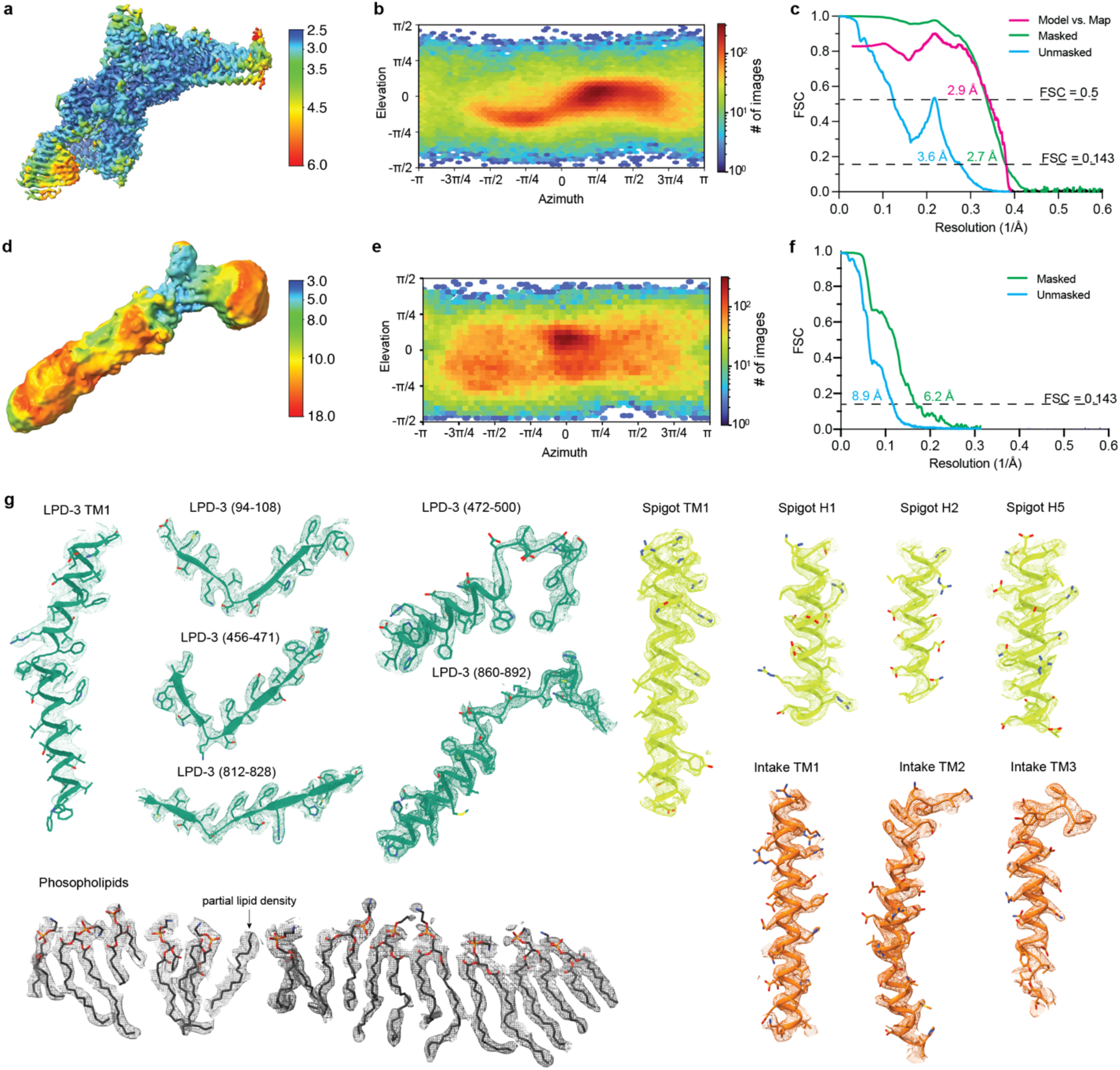
Cryo-EM statistics, angular distributions, and selected sections of density maps. **a,** N-terminal electron density map colored by local resolution values. **b,** Angular distributions of the final N-terminal reconstruction. **c,** Fourier shell correlations (FSEC) curves for the N-terminal map and model. **d,** Full-length electron density map colored by local resolution values. **e,** Angular distributions of the final full-length reconstruction. **f,** Fourier shell correlations (FSEC) curves for the full-length map and model. **g,** Fragments of cryo-EM density map and atomic model of LPD-3, Intake, Spigot, and phospholipid molecules in the tunnel. The cryo-EM maps are shown as mesh.

**Extended Data Figure 6.**
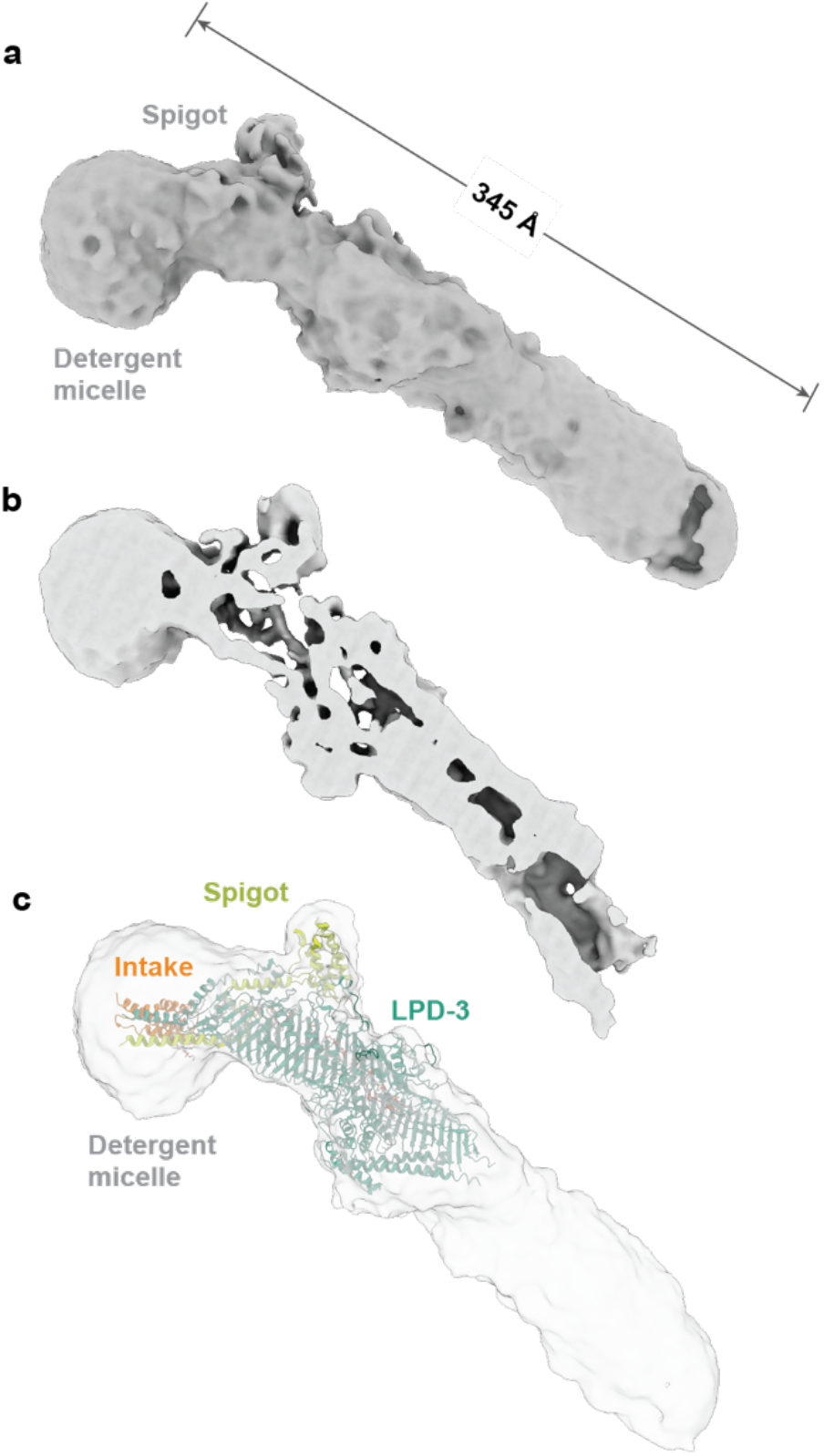
Cryo-EM density map of the full-length LPD-3 complex. **a,** Electron density map of the full-length LPD-3 complex, shown parallel to the membrane. **b,** Clipped view of the density map, showing that an internal cavity is observed along the entire length of the map. **c,** A model of the N-terminal LPD-3 complex is fit within the full-length density map. The full-length density map is shown in opaque grey and the model is colored as in Figure 1.

**Extended Data Figure 7.**
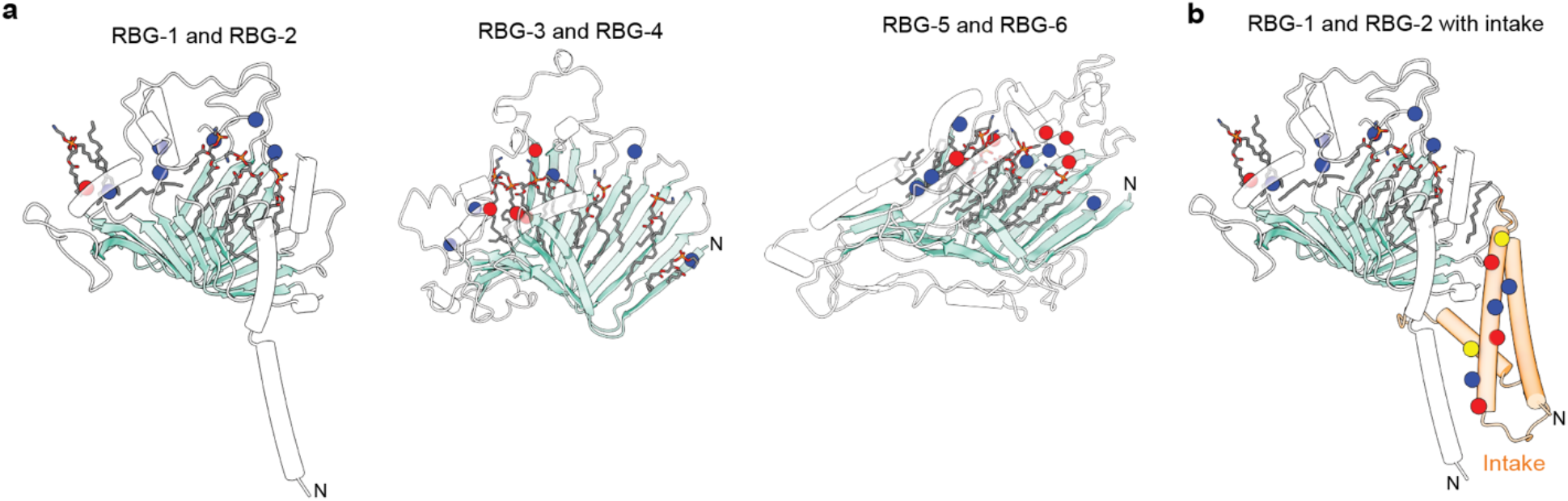
The lipid transport track. **a**, The locations of ionizable residues within the LPD-3 tunnel are shown as blue spheres for basic residues and red spheres for acidic residues. Lipids are shown grey sticks. The LPD-3 tunnel is divided into three sections for ease of viewing. The β-taco is colored light teal and the loop-cap and transmembrane helix are colored white. **b,** Intake is shown along with the LPD-3 RBG-1 and RBG-2 domains to further illustrate the continuous lipid transport track that extends from the ER lumen and through the tunnel. Intake is colored light orange and residues that interact with phospholipid headgroups in the MD simulations (Figure 4) are shown as spheres. Polar residues are shown as yellow spheres and other colors are the same as in (a).

**Table 1.**
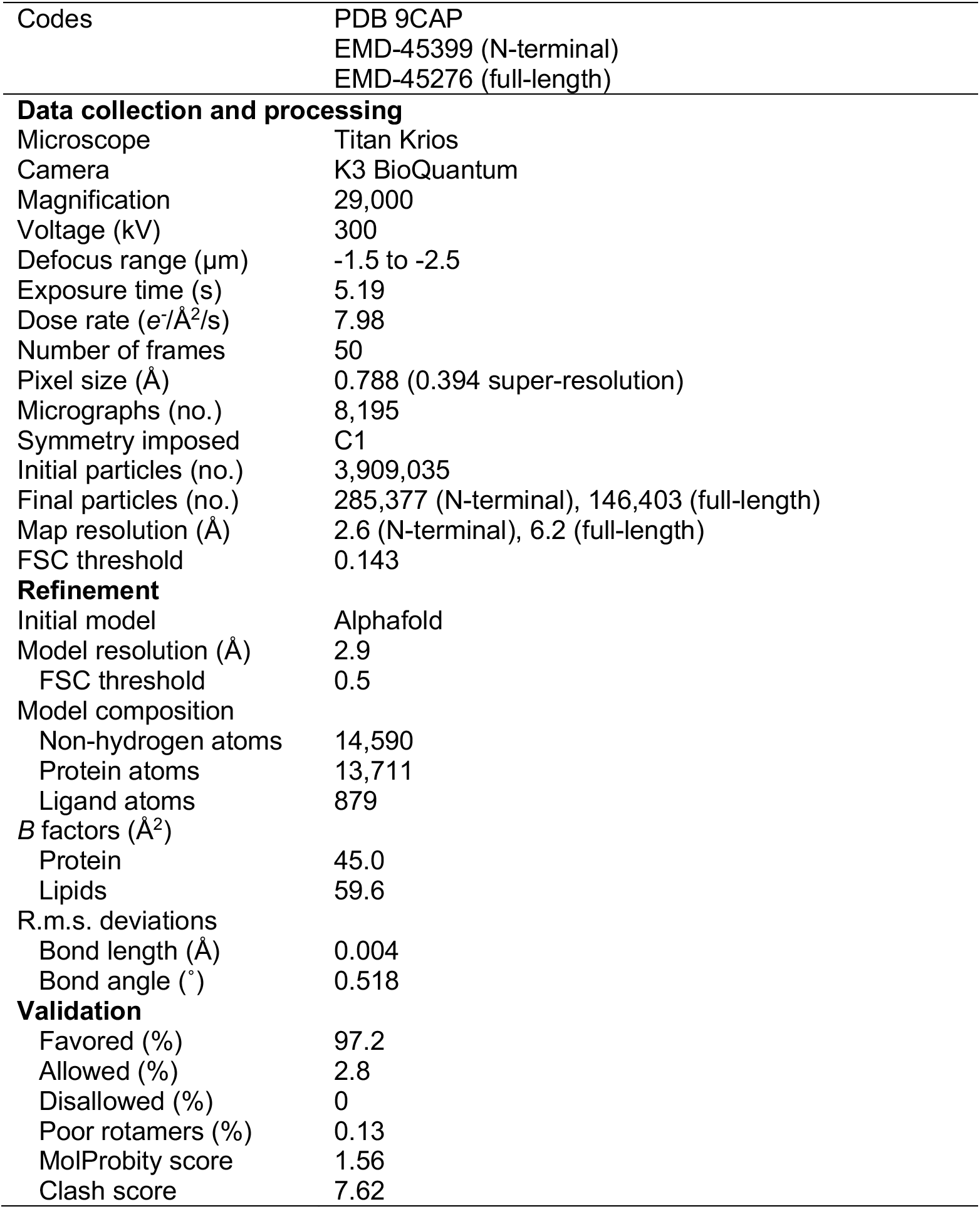
Cryo-EM statistics.

